# HyDrop: droplet-based scATAC-seq and scRNA-seq using dissolvable hydrogel beads

**DOI:** 10.1101/2021.06.04.447104

**Authors:** Florian V. De Rop, Joy N. Ismail, Carmen Bravo González-Blas, Gert J. Hulselmans, Christopher C. Flerin, Jasper Janssens, Koen Theunis, Valerie M. Christiaens, Jasper Wouters, Gabriele Marcassa, Joris de Wit, Suresh Poovathingal, Stein Aerts

## Abstract

Single-cell RNA-seq and single-cell ATAC-seq technologies are being used extensively to create cell type atlases for a wide range of organisms, tissues, and disease processes. To increase the scale of these atlases, lower the cost, and allow for more specialized multi-ome assays, custom droplet microfluidics may provide complementary solutions to commercial setups. We developed HyDrop, a flexible and generic droplet microfluidic platform encompassing three protocols. The first protocol involves creating dissolvable hydrogel beads with custom oligos that can be released in the droplets. In the second protocol, we demonstrate the use of these beads for HyDrop-ATAC, a low-cost non-commercial scATAC-seq protocol in droplets. After validating HyDrop-ATAC, we applied it to flash-frozen mouse cortex and generated 8,502 high-quality single-cell chromatin accessibility profiles in a single run. In the third protocol, we adapt both the reaction chemistry and the capture sequence of the barcoded hydrogel bead to capture mRNA, and demonstrate a significant improvement in throughput and sensitivity compared to previous open-source droplet-based scRNA-seq assays (Drop-seq and inDrop). Similarly, we applied HyDrop-RNA to flash-frozen mouse cortex and generated 9,508 single-cell transcriptomes closely matching reference single-cell gene expression data. Finally, we leveraged HyDrop-RNA’s high capture rate to analyse a small population of FAC-sorted neurons from the *Drosophila* brain, confirming the protocol’s applicability to low-input samples and small cells. HyDrop is currently capable of generating single-cell data in high throughput and at a reduced cost compared to commercial methods, and we envision that HyDrop can be further developed to be compatible with novel (multi-) omics protocols.

## Main

Droplet-microfluidic single-cell sequencing technologies have enabled the profiling of tens of thousands^1,2^ - and recently millions^3,4^ - of single cells. Owing to their limited sensitivity (e.g. median of 250-500 genes per cell in primary tissues^1,5^), and relatively lengthy workflows compared to commercial solutions such as 10x Genomics, generic protocols such as Drop-seq^6^ and InDrop^7^ have been used much less than the commercial alternatives ^5,8,9^. A second wave of droplet-based assays has provided the ability to profile chromatin accessibility of single cells, particularly using single-cell ATAC-seq^10^. To our knowledge, only one non-commercial droplet-based scATAC-seq protocol has been published so far^11^. Despite its elegant conceptual solution to droplet-based scATAC/scRNA-seq, the SNARE-seq protocol is labour intensive compared to commercial solutions such as 10x Genomics’ Chromium^12^ and Biorad’s ddSEQ^13^, and the use of resin beads in the SNARE-seq protocol leads to reduced cell capture and sensitivity. We developed HyDrop, a new hydrogel-based droplet microfluidic platform to improve the sensitivity and usability of both scRNA-seq and scATAC-seq in droplets, and to provide the first hydrogel-based solution for droplet-based scATAC-seq. All HyDrop protocols and analysis code are freely available at https://www.protocols.io/workspaces/aertslab and https://hydrop.aertslab.org.

In the first HyDrop protocol, we generate barcoded hydrogel beads that can dissolve and release their embedded barcoded oligonucleotide. Polyacrylamide beads incorporating disulfide crosslinkers and short oligonucleotide PCR handles are generated by droplet microfluidics similar to a recently published method^14^. A custom droplet microfluidic chip (fig. S1) is employed to produce beads of approximately 45 μm diameter, but flow rates can be tuned to change bead diameter. These hydrogel beads are then barcoded using a modified three-round split-pool PCR synthesis method^7,15^, resulting in 96×96×96 (884,736) barcode possibilities. The terminal sequence used in the final round of barcoding can be varied depending on the assay the beads will be used for (see Methods, fig. 1a). A sequence complementary to the Tn5 transposase adapter is used to capture tagmented chromatin fragments in scATAC-seq and a unique molecular identifiers (UMI)^16^ + poly(dT) sequence is used to capture and count poly(A)+ mRNA in scRNA-seq (see further below under the HyDrop-RNA protocol). The barcoded beads are stored in a glycerol-based freezing buffer at −80 °C in order to prevent loss of primers over time (fig. S2a).

**Figure 1.**
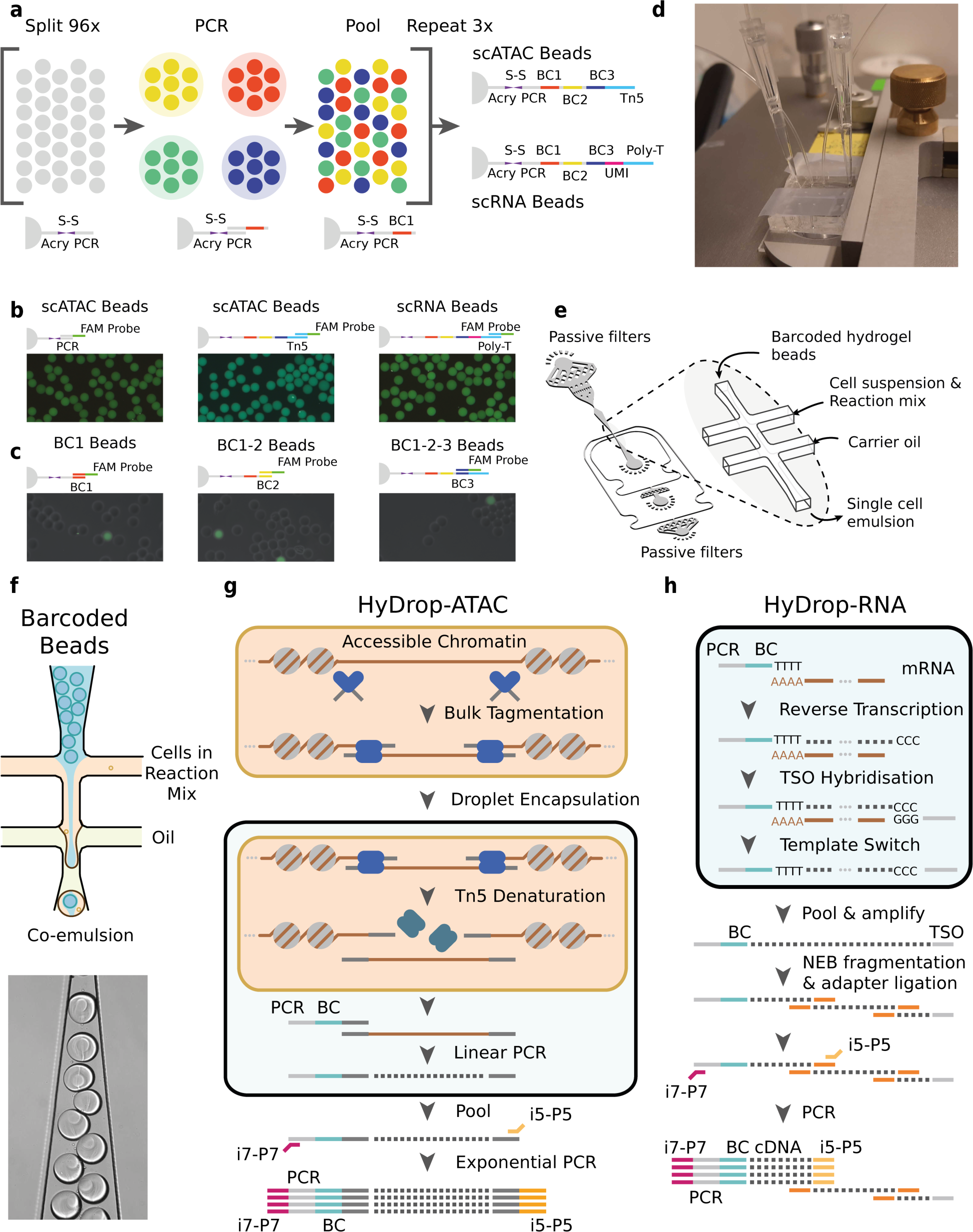
**a**. Split-pool process for barcoding of dissolvable hydrogel beads. Beads are sequentially distributed over 96 wells, sub-barcoded, re-pooled, and distributed three times to generate 96×96×96 (884736) possible barcode combinations. Different 3’-terminal capture sequences are possible depending on the oligonucleotide sequence appended in the last step. **b**. Semi-quantitative assessment of bead primer incorporation by FISH after every sub-barcoding step shows that bead fluorescence uniformity is retained throughout the barcoding process. **c**. FISH with probes complementary to only one of 96 sub-barcode possibilities shows that approximately 1/96 beads exhibit fluorescence for a selected sub-barcode probe. Fluorescence signal is overlaid with a brightfield image at 50% transparency to indicate positions of non-fluorescent beads. **d**. Microfluidic chip setup on the Onyx platform. Cells and beads are loaded into pipette tips and plugged into a HyDrop Chip. Flow of oil and aqueous phases is achieved by Onyx displacement syringe pumps. **e**. HyDrop chip design has three inlets: one each for carrier oil, barcoded hydrogel beads and cell/reaction mix. Passive filters at each inlet prevent dust and debris from entering the droplet generating junction. **f**. Diagram and snapshot of cell/bead droplet encapsulation. Schematic overview of HyDrop-ATAC **(g)** and HyDrop-RNA **(h)** assay for single-cell library generation. Nuclear membrane is visualised in salmon, water droplet is visualised in blue.

We validated the purity and concentration of the hydrogel bead primers using fluorescent probes complementary to the beads 3-prime terminal sequence^15^ (fig. 1b) or one of the 96 sub-barcode possibilities (fig. 1c). These experiments show that there is no significant loss of primers or mixing of barcodes throughout the barcoding process, and that the beads are uniform in size and primer content. Additional testing revealed that our modified PCR barcoding method produced more uniformly barcoded beads compared to the isothermal amplification protocol described in inDrop (fig. S2b). Furthermore, the disulfide moieties incorporated in both the bead’s polymer matrix and oligonucleotide linker can be cleaved when exposed to reducing conditions, such as DTT. This chemical method of release is more user-friendly compared to the UV-mediated^7,15^ primer release as the beads do not have to be shielded from light. Furthermore, the entire barcoding reaction can be executed in the reducing environment standard to many biochemical reactions. In addition to improved primer release compared to non-dissolvable beads (fig. S2c), dissolvable beads also do not disrupt the emulsion during the thermocycling needed for scATAC-seq (see methods) as they dissolve within minutes in the droplet’s reducing environment (fig. S3a, b). Finally, by varying the concentration of the acrydite primer during bead synthesis, lower or higher amounts of cleavable barcoded primers can be generated. When the acrydite primer concentration incorporated in the bead is high (50 μM, similar to InDrop^15^), excess unreacted barcodes cannot be sufficiently filtered out in further downstream steps. These primers are carried over to subsequent reactions, leading to random barcoding of free fragments after droplet merging, and subsequently to cell-mixed expression or chromatin accessibility profiles. The bead primer concentration with an optimal balance between sensitivity and library purity was found to be 6 to 12 µM for both scATAC-seq and scRNA-seq (fig. S4a, b).

Our second HyDrop protocol provides a generic assay for droplet-based scATAC-seq. Here, nuclei are co-encapsulated with HyDrop-ATAC beads after tagmentation in bulk. In order to co-encapsulate beads and nuclei with a high capture rate and minimal microfluidic complexity, we developed a custom microfluidic chip (fig. S1). The chip design features two inlets for beads and cells or nuclei and one inlet for the emulsion carrier oil. Several layers of passive filters near the inlet ports mitigate dust and debris buildup during droplet generation to prevent obstruction of the channels. Beads and nuclei are loaded via a tip reservoir to reduce non-linear flow behaviour and the potential accumulation of cells/nuclei and hydrogel beads associated with narrow tubes^17,18^ (fig. 1d, e). Due to the stability of all flows and the deformable nature of the hydrogel beads, > 90% occupancy of hydrogel beads in droplets can be achieved^19^ (fig. 1f). After co-encapsulation with the tagmented nuclei, the hydrogel beads dissolve in the presence of DTT in the nuclei/PCR mix and release their uniquely barcoded primers inside the droplet as described above. Subsequent thermocycling of the emulsion denatures the Tn5 protein complex and releases accessible chromatin fragments within the droplet. These fragments are then linearly amplified and cell-indexed by the bead’s barcoded primers after which the emulsion is broken and the indexed ATAC fragments are pooled, PCR amplified, and sequenced (fig. 1g). Pitstop, a selective small molecule inhibitor of clathrin is supplemented during nuclei extraction and tagmentation to increase nucleus permeability to Tn5^20^.

To assess the purity of scATAC-seq libraries generated by HyDrop-ATAC, we performed two mixed-species experiments. First, we generated single-nuclei ATAC-seq libraries from a 50:50 mixture of human breast cancer (MCF-7) and a mouse melanoma cell line generated previously^21^. For the pre-processing and mapping of HyDrop-ATAC data, we developed a custom preprocessing pipeline^22^. After filtering the cell barcodes for a minimum TSS enrichment score of 7 and unique fragment count of 1,000, we recovered 1,353 cells from a target of 2,000, with a median of 2,705 unique fragments per cell. We identified 98.4% of cells as either human or mouse at a minimum purity of 95% fragments mapping to either species (fig. 2a). This implied doublet rate of 3.1% is comparable to other droplet microfluidic protocols^6,7,12^. Next, we generated libraries from a mixture of MCF-7/PC-3/Mouse cortex (45:45:10) to evaluate whether two human cell types can be distinguished. A spike-in of 10% mouse cells was used as an internal control. We recovered 2,602 human cells, 466 mouse cells, and 93 species doublets after filtering for 95% species purity. Clustering human cells (together with the MCF-7 cells from the first species mixing experiment to evaluate batch effects) recovered two distinct populations, each exhibiting specific ATAC-seq peaks near MCF-7 or PC-3 marker genes (fig. S5a). Aggregated reads per cluster showed typical ATAC-seq profiles concordant with public bulk ATAC-seq data^23^ (fig. 2b, fig. S5b).

**Figure 2.**
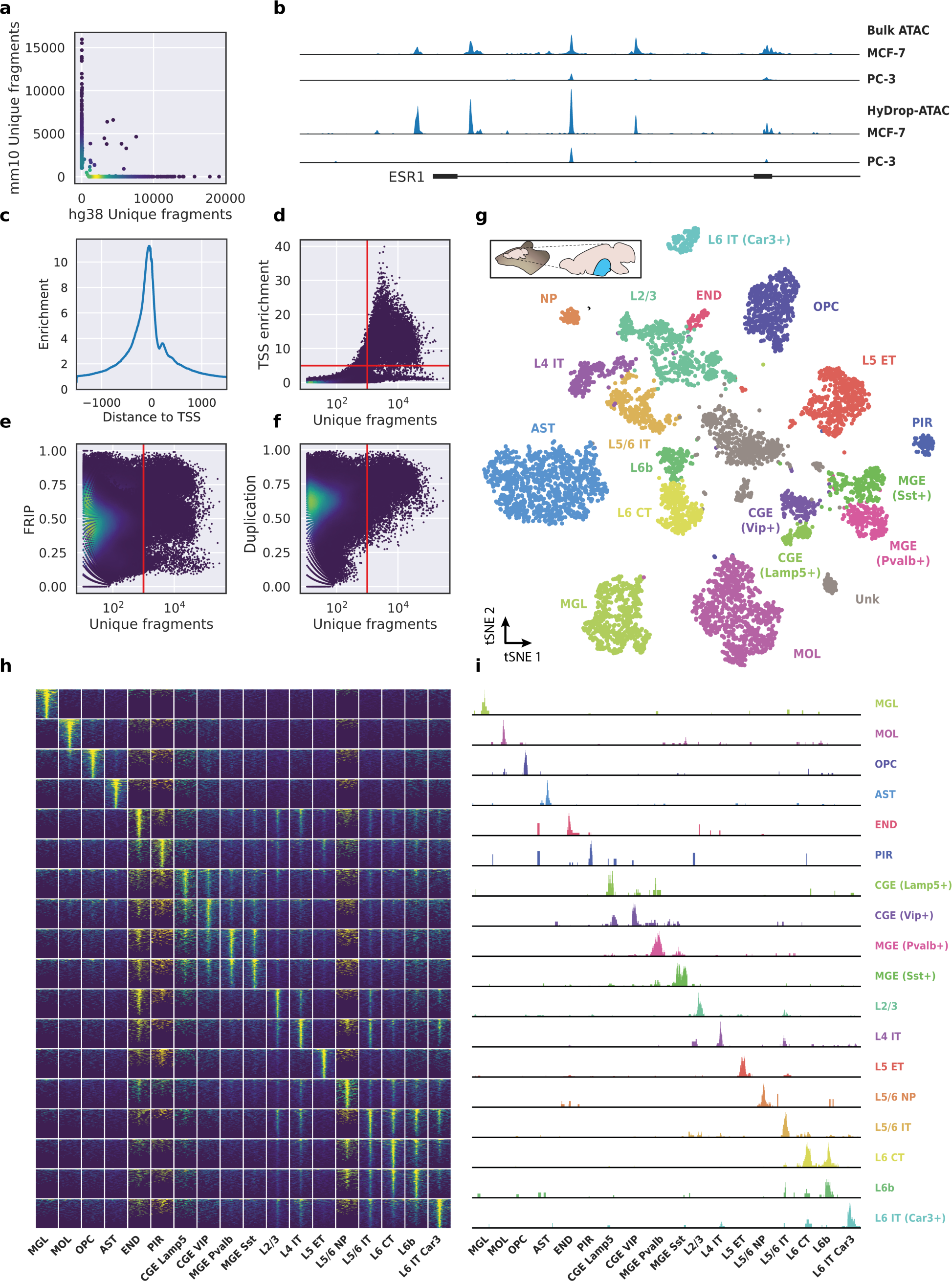
**a**. Scatterplot of the number of unique fragments detected in a 50:50 mixture of human MCF-7 and mouse melanoma cells coloured by local density estimation. **b**. RPGC-normalized aggregate genome tracks comparing HyDrop-ATAC and bulk ATAC-seq profiles of human MCF-7 and PC-3 cell lines around the Estrogen receptor 1 (ESR1) locus, scaled to maximum of all four samples. Aggregate enrichment profile of reads around transcription start site (TSS) **(c)**, TSS enrichment per barcode **(d)**, fraction of reads in peaks (FRIP) per barcode **(e)** and duplication rate per barcode **(f)** in mouse cortex HyDrop-ATAC data. A minimum TSS enrichment of 5 and a unique number of fragments of 1000 are used as cut-off values to separate cells from background (red lines). Cells are colored by local density estimation. **g**. UMAP projection of 8502 mouse cortex nuclei annotated with cell type inferred by accessibility near marker genes. Abbreviations: microglia (MGL), mature oligodendrocytes (MOL), oligodendrocyte precursors (OPC), astrocytes (AST), endothelial cells (END), piriform cortex neurons (PIR), caudal and medial ganglionic eminence derived neurons (CGE, MGE), layers 2-6 intratelencephalic (IT), L5 extratelencephalic (ET), L5/6 near projecting excitatory neurons (NP), L6 corticoencephalic (CT), and deep L6 excitatory neurons (L6b). **h**. Aggregate accessibility of top 1000 differentially accessible regions per cluster. **i**. Row-scaled, CPM-normalized aggregate genome track covering the top 1 differentially accessible region (DAR) for each cluster.

To evaluate the performance of HyDrop-ATAC on primary tissue, we then generated single cell libraries from snap-frozen, dissected adult mouse brain cortex. Libraries were sequenced to approximately 75% duplication rate. After filtering for a minimum of 1,000 unique nuclear fragments, a TSS enrichment score of 5, and removing 481 cells (5.4%) detected as doublets by Scrublet^24^, we recovered a total of 8,502 single nuclei. Cells passing the filters had a median of 4,148 fragments per cell, a median TSS enrichment score of 13, and a median of 53% of fragments in peaks, reflecting high-quality cells and low levels of background signal (fig. 2c-f). Even though the number of unique fragments per cell (~4K) is lower than that of commercial methods (e.g., 17-20K per cell for 10x Genomics, see Methods), HyDrop-ATAC compares favourably with these platforms in terms of TSS enrichment and FRIP scores, and due to the possibility to profile large cell numbers (>8K cells in a single run), cell type clustering of mouse brain achieved higher resolution compared to publicly available 10x Genomics data sets. We used cisTopic^25^ to reduce the dimensionality of the dataset and the Leiden algorithm^26^ to cluster cells (fig. 2g). Cell annotation using the aggregated ATAC signal around several neocortex markers^27,28^ recovered 19 distinct cell types, similar to previously published scATAC-seq mouse cortex data^29,30^ (fig. S6). For example, we identified oligodendrocyte precursors and mature oligodendrocytes, marked by exclusive accessibility nearby *Sox10* and *Pdgfra2*, respectively. Within ganglionic eminence-derived interneurons, we were able to further distinguish medial ganglionic eminence-derived subtypes with specific ATAC-seq signal near either *Vip* or *Lamp5*, and caudal ganglionic eminence-derived subtypes with accessibility near either *Sst* or *Pvalb*. Finally, HyDrop-ATAC data revealed distinct cell-type specific differentially accessible regions (fig. 2h-i).

In the third HyDrop protocol, we implemented a new scRNA-seq assay using barcoded bead primers carrying a 3-prime poly(dT) sequence. Single cells or nuclei are resuspended in a reverse transcriptase mix and co-encapsulated into microdroplets with 3-prime poly(dT) HyDrop beads. The same microfluidic chip design is used for both HyDrop-RNA and HyDrop-ATAC. Cells are lysed inside the droplets upon contact with the lysis buffer in which the barcoded beads are suspended. Simultaneously, barcoded primers are released from the hydrogel bead after exposure to DTT present in the reverse transcriptase mix. Reverse transcription inside the emulsion generates thousands of barcoded single-cell cDNA libraries in parallel. The emulsion is then broken and the single-cell transcriptome libraries are processed further in a pooled manner (fig. 1h), similarly to the InDrop protocol^15^. However, although both assays are based on hydrogel beads, HyDrop-RNA differs significantly from InDrop. HyDrop-RNA employs a template switching oligo (TSO) reverse transcription technique (similar to Drop-seq), rather than an *in vitro* transcription/random hexamer priming workflow. This change simplifies and speeds up the protocol significantly with no reduction in sensitivity. To optimize the sensitivity of the assay, we compared several different reaction chemistries: (1) Exonuclease I treatment to remove excess of unused barcode primers; (2) the use of a locked nucleic acid (LNA) TSO^31^; (3) the addition of GTP/PEG into the reverse transcription step (similar to SMART-seq3^32^); and (4) the use of second strand synthesis. For the latter, we tested both alkaline hydrolysis and enzymatic treatment (RNAse H) to remove the RNA strand from the first strand product, and we evaluated the performance of both the Bst 2.0 DNA polymerase and the Klenow (exo-)^33,34^ fragment for second strand synthesis. By comparing these variations on a 50:50 human-mouse (human melanoma, mouse melanoma) mixture, we found that the GTP/PEG protocol with Exonuclease I treatment performed best, yielding a median of 2,110 UMIs and 1,325 genes per cell with a species-purity of 90.1% (fig. 3a, S7). Accordingly, the GTP/PEG method was used in all further HyDrop-RNA experiments. Applying this protocol to a 50:50 human-mouse (MCF-7, mouse melanoma) mixture recovered 1,235 human and 846 mouse cells with 169 species doublets at a cutoff of 95% species purity, with a median of 1,439 UMIs and 904 genes per cell (fig. 3b).

**Figure 3.**
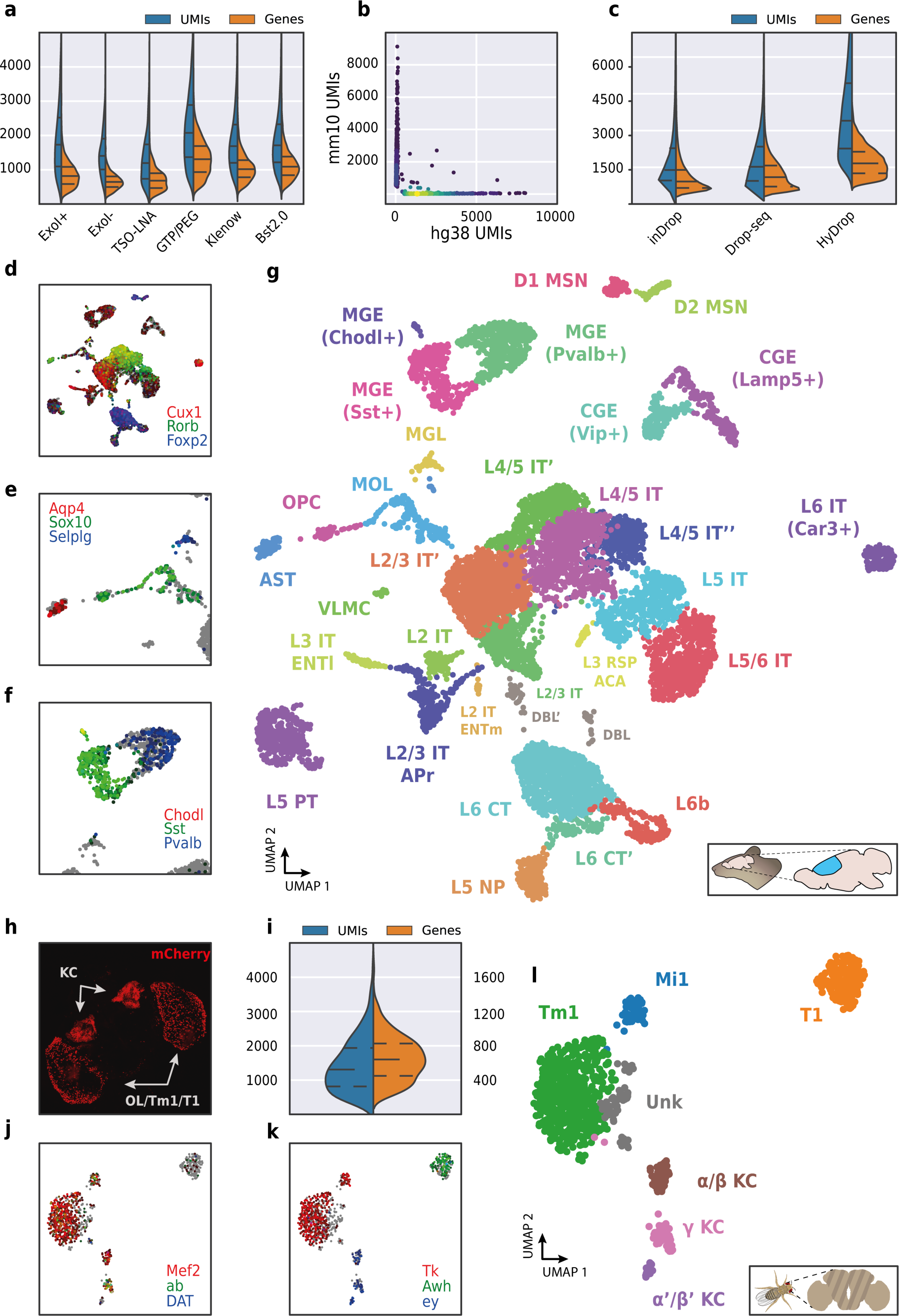
**a**. Comparison of UMI and gene count of HyDrop-RNA with and without Exo I treatment post-droplet merging, with the use of a locked nucleic acid (LNA) template switching oligo (TSO) and with GTP/PEG, BST2.0 and Klenow fragment library preparation. Inner lines represent Q1, median and Q3. **b**. Scatterplot of human and mouse UMIs detected in a 50:50 mixture of human MCF-7 and mouse melanoma cells coloured by local density estimation. **c**. Comparison of UMI and gene count of public inDrop mouse cortex data, public Drop-seq mouse retina data, and HyDrop-RNA mouse cortex data. Inner lines represent Q1, median and Q3. Mouse cortex UMAP is colored by log-scaled UMI counts of Cux1, Rorb, Foxp2 **(d)**, Aqp4, Sox10, Selplg **(e)**, Chodl, Sst and Pvalb **(f)**. Colors are scaled to minimum and maximum values. **g**. UMAP projection of 9507 mouse cortex nuclei annotated with cell type inferred by marker gene expression. Abbreviations: microglia (MGL), mature oligodendrocytes (MOL), oligodendrocyte precursors (OPC), astrocytes (AST), endothelial cells (END), piriform cortex neurons (PIR), caudal and medial ganglionic eminence derived neurons (CGE, MGE), layers 2-6 intratelencephalic (IT), pyramidal tract (PT), near projecting excitatory neurons (NP) and corticoencephalic (CT) neurons, layer 2 intratelencephalic medial entorhinal area neurons (L2 IT ENTm), L2/3 intratelencephalic area prostriata neurons (L2/3 IT APr), layer 3 intratelencephalic entorhinal neurons (L3 IT ENTl), layer retrosplenial and anterior cingulate area neurons (L3 RSP ACA), deep L6 excitatory neurons (L6b), D1 and D2 medium spiny neurons (MSN), and vascular leptomeningeal cells (VLMC). **h**. Confocal maximum intensity projection of R74G01-Gal4>UAS-mCherry brain. **i**. Violin plot of UMIs and genes detected in nuclei derived from FAC-sorted fly neurons. Inner lines represent Q1, median and Q3. Fly neuron UMAP colored by log-scaled UMI counts of Mef2, ab, DAT **(j)** and Tk, Awh, ey **(k)**. Colors are scaled to minimum and maximum values. **l**. UMAP of 973 FAC-sorted *Drosophila* neurons annotated with cell types inferred by marker gene expression (KC, Kenyon cells; Tm1, transmedullary neuron; Mi1, medullary intrinsic neuron).

We then used HyDrop-RNA to generate 9,508 single nuclei transcriptomes from snap-frozen mouse cortex in a single experiment. At a saturation of approximately 60% duplicates and with > 55% of reads mapping to transcriptome, we achieve a median of 3,404 UMIs and 1,662 genes per cell after filtering (fig. 2c), a significant improvement over the median of 1,521 UMIs and 1,097 genes reported by inDrop snRNA-seq on mouse auditory cortex neurons^5^ and the median of 1,389 UMIs and 922 genes reported by Drop-seq on mouse retina neurons^6^. 10x Chromium v2 gene expression reference data reports a median genes of 775-2,679 and a median UMIs of 1,127-6,570 on E18 and adult mouse brain nuclei (see methods). Comparing the top per-cluster differentially expressed genes with markers from the Allen Brain Atlas SMART-seq data^27^ revealed 30 distinct populations corresponding to previously identified cell types (fig. 2d-f, fig. S8). In addition to the major neuronal and glial populations previously detected in our HyDrop-ATAC experiment, we detect a small population of vascular leptomeningeal cells (VLMC) and layer 2 intratelencephalic neurons from the medial entorhinal area (L2 IT ENTm). We also detect both D1 and D2 medium spiny neurons (MSN) as a result of residual striatal tissue and layer 3 *Scnn1a*+ neurons from the retrosplenial and anterior cingulate area (L3 RSP ACA), concordant with Atlas SMART-seq data^27^.

To assess HyDrop-RNA’s performance on low cell input samples, we performed the protocol on approximately 1500 FAC-sorted neurons from the *Drosophila* brain. We dissected brains in which mCherry expression was driven in specific cell populations by a Gal4 driver line (R74G01-Gal4) and used mCherry-positive cells as input for HyDrop-RNA (fig. 2h). Of the 1,500 cells obtained after FACS sorting, we recovered 973 fly brain cells with a median of 1,307 UMIs and 640 genes (fig. 2i). In-house Drop-seq performed on fly brain neurons recovered a median of 579 UMIs and 289 genes per cell at a deeper sequencing saturation^1^. Annotation of the 973 single-cell transcriptomes obtained by HyDrop confirmed the presence of all three Kenyon cell subtypes alongside T1 and Tm1 neurons, as expected from our stainings and previous reports^35^. Surprisingly, we also detected a small population of Mi1 neurons (fig. j-l, fig. S9).

By applying HyDrop to generate thousands of mouse, human, and Drosophila single-cell gene expression and chromatin accessibility profiles, we demonstrate the protocol’s applicability to a variety of different biological samples. Our experiments on mouse and fly tissues recapitulate cellular heterogeneity in these complex samples and strongly agree with reference datasets from both organisms. We show that HyDrop outperforms its open-source predecessors in terms of sensitivity and user-friendliness. Moreover, at a per-cell library cost of < $0.03 it does so at a significantly lower cost compared to commercial droplet-microfluidic alternatives. We envision that this reduction in both cost and labour will accelerate the scaling of large-scale atlasing efforts and bring the benefits of single-cell sequencing to smaller projects. We believe that further optimization and modification of the protocol’s reaction chemistry and bead composition will lead to improvements in sensitivity and stimulate the development of novel (multi-) omics droplet-based assays. Additionally, HyDrop’s flexible hydrogel bead synthesis toolkit may potentially be exploited to design more complex single-cell (multi-) omics assays such as the capture of accessible chromatin, (m)RNA, and proteins or antibodies from the same single cell.

## Methods

### Microfluidic droplet generator manufacturing

Microfluidic droplet generators were produced using standard SU-8 lithography and PDMS lithography according to well established protocols^36^. Briefly, the design for droplet generators were made in AutoCAD R2014 and the designs are printed onto a chrome mask using a laser writer. The SU-8 lithography is performed on a 4 inch silicon wafer using SU 8-2050 (Microchem) negative photoresist using UV aligner (EVG-620). As per manufacturers’ recommendation, spin coating of the wafer with SU8 was performed at 500 rpm (ramp 100rpm/s) −10s and 2000 rpm (ramp: 300rpm/s) −30s, to achieve a feature height of 70-80 um). For preparing the PDMS chip, a mixture of PDMS monomer and crosslinker (Dow Corning SYLGARD 184) was prepared at a ratio of 10:1 and mixed thoroughly. The mixture was degassed in vacuum for 45 minutes and poured onto an SU-8 master template and baked at 80 °C for 4 hours. Inlet ports were cut using a 1 mm biopsy needle after which the chips were exposed to high-voltage plasma for 30 s and bonded onto a glass slide. 5 μL of 2% Trichloro(1H,1H,2H,2H-perfluorooctyl)silane in HFE was injected into each channel, incubated for 10 mins at room temperature and excess oil was removed by applying pressurized air. Chips were finally baked at 100 °C for 2 hours (more detailed methods for photolithography and PDMS lithography in https://www.protocols.io/workspaces/aertslab).

### Barcoded hydrogel bead manufacturing & storage

Dissolvable hydrogel beads are synthesized similar to a previously published protocol^14^ and barcoded according to a modified inDrop protocol^15^. For synthesizing 2-3 mL batch of beads, 2 mL of Bead Monomer Mix (6% acrylamide, 0.55% bisacryloylcystoylamine, 10% TBSET (10 mM

Tris-HCl pH 8, 137 mM NaCl, 2.7 mM KCl, 10 mM EDTA, 0.1% Triton X-100), 12 μM acrydite primer, 0.6% ammonium persulfate) was encapsulated into 50 μm diameter droplets in HFE-7500 Novac oil with EA-008 surfactant (RAN Biotech). 1 mL aliquots of the resulting emulsion was layered with 400 μL of mineral oil and incubated at 65 °C for 14 hours. Excess mineral oil and the emulsion oil was removed and 2-3 washes with 1 mL of droplet breaking solution (20% PFO in HFE) was performed. Beads were pelleted at 5000 xg, 4 °C for 30 seconds and washed twice in 1 mL of 1% SPAN-80 in hexane. Beads were sequentially washed in TBSET until all hexane phase was removed.

Beads were washed twice in Bead Wash Buffer (10 mM Tris-HCl pH 8, 0.1% Tween-20), twice in PCR Wash Buffer (10 mM Tris-HCl pH8, 50 mM KCl, 1.5 mM MgCl2, 0.1% Tween-20). The subsequent liquid handling in the 96-well plate is performed using Hamilton microlab STAR robot. 22.5 μL of beads were distributed to a 96 well plate. 2.5 μL of 100 μM sub-barcode primer and 25 μL of KAPA HiFi Hotstart master mix (Roche) was added to each well and the plate was thermocycled (95 °C 3 min., 5 cycles of 98 °C 20s, 38 °C 4 min., 72 °C 2 min., 1 cycle of 98 °C 1 min., 38 °C 10 min., 72 °C 4 min., followed by a final hold on 4 °C) with intermittent vortexing during every annealing step. 50 μL of STOP-25 (10 mM Tris-HCl pH 8, 25 mM EDTA, 0.1% Tween-20, 100 mM KCl) was added to each well to deactivate the polymerase and its contents were pooled. Remaining beads in wells were washed out with 100 μL of STOP-25 and the beads were rotated at room temperature for 30 minutes. Beads were then washed with STOP-10 (10 mM Tris-HCl pH 8, 10 mM EDTA, 0.1% Tween-20, 100 mM KCl) and rotated for 10 minutes in Denaturation Solution (150 mM NaOH, 85 mM BRIJ-35). Beads were washed twice in Denaturation Solution and twice more in Neutralisation Solution (100 mM Tris-HCl pH 8, 10 mM EDTA, 0.1% Tween-20, 100 mM NaCl). The sub-barcoding step was repeated twice more for a total of 3 sub-barcodes.

Hydrogel beads were sequentially filtered using a 70 μm strainer (Falcon). For both the HyDrop-ATAC and RNA beads were pelleted at 300 xg, 4 °C and resuspended in 5 mL of Bead Freezing Buffer (150 mM NaCl, 125 mM Tris-HCl pH 7, 10 mM MgCl2, 4% Tween-20, 0.75% Triton X-100, 30% glycerol, 0.3% BSA). Beads were pelleted at 300 xg, 4 °C and resuspended in 5 mL of Bead Freezing Buffer a second time and incubated at 4 °C for at least 3 hours. Beads were pelleted at 1000 xg, 4 °C and the pellet was aliquoted into 35 μL aliquots and stored at −80 °C for long term storage (further method details in https://www.protocols.io/workspaces/aertslab)

### Hydrogel bead fluorescence in-situ hybridisation quality control

Bead QC was performed as described previously^15^. Briefly, 10 μL of hydrogel beads were resuspended in 1 mL of hybridization buffer (5 mM Tris-HCl pH 8.0, 5 mM EDTA, 0.05% Tween-20, 1M KCl) and centrifuged for 1 min. at 1000 xg. The wash was repeated once more, and 960 μL of the supernatant was removed. 4 μL of 200 μM specific FAM probe was added depending on which part of the barcode needed testing (see supplementary table 1). The beads were incubated at room temperature in the dark for 30 min. Beads were washed thrice in QC buffer and visualised under a Zeiss Axioplan 2 microscope (300 ms exposure time, 80% lamp intensity).

### Cell culture and cell dissociation

MCF-7 cells were cultured in RPMI1640 (ThermoFisher 11875093) medium supplemented with 10% FBS (ThermoFisher 10270-106), 1% penicillin/streptomycin (Life Technologies 15140122), and 10 ug/mL insulin (Sigma Aldrich I9278) and passaged twice per week. PC-3 cells were cultured in RPMI1640 medium supplemented with 10% FBS and 1% penicillin/streptomycin and passaged twice per week. Mouse melanoma cells were cultured in DMEM (ThermoFisher 13345364) supplemented with 10% FBS and 1% penicillin/streptomycin and passaged once per week. MM087 melanoma cells were cultured in F-10 Nutrient mix supplemented with 10% FBS and 1% penicillin/streptomycin and passaged once per week. Cells were washed in PBS and dissociated into single cell suspensions by adding 1.5 mL of 0.05% Trypsin (Life Technologies 25300054) and waiting for 5 to 7 minutes. The single-cell suspension was centrifuged at 500 rcf for 5 min at 4°C and the resulting pellet was resuspended in PBS. This PBS wash was repeated once more and the single-cell suspension was processed further.

### Fly rearing and cell dissociation

GMR74G01-Gal4 (BL#39868) and UAS-mCherry (BL#38425) flies were obtained from Bloomington Drosophila Stock Center. The resulting cross strain was raised on standard cornmeal-agar medium at 25°C at a 12h light/dark cycle. 50 adult brains were dissected in DPBS and transferred to a tube containing 100 μL of cold DPBS solution. Samples were centrifuged at 500 rcf for 1 min and the supernatant was replaced by 50 μL of dispase (3 mg/mL, Sigma-Aldrich, D4818, 2mg) and 75 μL of collagenase I (100 mg/mL, Invitrogen, 17100-017). Brains were dissociated in a Thermoshaker (Grant Bio PCMT) at 500 rpm for 2 h at 25°C, with pipette mixing every 15 min. Cells were subsequently washed with 1000 μL of cold DPBS solution and centrifuged at 500 rcf for 5 min at 4°C and resuspended in 250 μL of DPBS with 0.04% BSA. Cell suspensions were passed through a 10 μM pluriStrainer (ImTec Diagnostics, 435001050). Cells were sorted based on viability and mCherry positivity using the Sony MA900 cell sorter. Sorted cells were collected into Eppendorf tubes pre-coated with 1% BSA.

### Cell line nuclei extraction

A pellet of 1 million dissociated cells or fewer was incubated on ice in 200 μL of ATAC Lysis Buffer (1% BSA, 10 mM Tris-HCl pH 7.5, 10 mM NaCl, 0.1% Tween-20, 0.1% NP-40, 3 mM MgCl2, 70 μM Pitstop, 0.01% Digitonin) for 5 to 7 minutes. 1 mL of ATAC Nuclei Wash Buffer (1% BSA, 10 mM Tris-HCl pH 7.5, 0.1% Tween-20, 10 mM NaCl, 3 mM MgCl2) was added and the nuclei were pelleted at 500 xg, 4 °C for 5 minutes. The resulting pellet was resuspended in 100 μL of ice-cold PBS and filtered with a 40 μm strainer (Flowmi).

### Mouse cortex dissection

All animal experiments were conducted according to the KU Leuven ethical guidelines and approved by the KU Leuven Ethical Committee for Animal Experimentation (approved protocol numbers ECD P183/2017). Mice were maintained in a specific pathogen-free facility under standard housing conditions with continuous access to food and water. Mice used in the study were 57 days old and were maintained on 14 h light, 10 h dark light cycle from 7 to 21 hours. In this study, cortical brain tissue from female P57 BL/6Jax was used. Animals were anesthetized with isoflurane, and decapitated. Cortices were collected, divided in four equal quadrants along the dorso-ventral and anterior-posterior axis, and immediately snap-frozen in liquid nitrogen. For HyDrop-ATAC, a ventral/posterior quadrant from the left hemisphere was used. For HyDrop-RNA, a dorsal/anterior quadrant was used from the left hemisphere of a second mouse.

### Snap-frozen mouse cortex nuclei extraction

For the preparation of nuclei for RNAseq, we used a modified protocol from the recently published single nuclei preparation toolbox^37^ to extract nuclei from snap-frozen mouse cortex samples. Briefly, a ~1 cm^3^ frozen piece of mouse cortex tissue was transferred to 0.5 mL of ice-cold homogenisation buffer (Salt-tris solution - 146 mM NaCl, 10 mM Tris 7.5, 1 mM CaCl2, 21mM MgCl2, 250 mM Sucrose, 0.03% Tween-20, 0.01% BSA, 25 mM KCl, 1mM 2-Mercaptoethanol, 1X cOmplete protease inhibitor, 0.5U/ul of RNAse In Plus (Promega)) in a Dounce homogenizer mortar and thawed for 2minutes. The tissue was homogenised with 10 strokes of pestle A and 5 strokes of pestle B until a homogeneous nuclei suspension was achieved. The resulting homogenate was filtered through a 70 μm cell strainer (Corning). The homogenizer and the filter is rinsed with an additional 500 ul of homogenization buffer. The tissue material was pelleted at 500xg and the supernatant was discarded. The tissue pellet was resuspended in a homogenization buffer without Tween-20. An addition 1.65 ml of homogenization buffer was topped up and mixed with 2.65 ml of Gradient Medium (75 mM sucrose, 1mM CaCl2, 50% Optiprep, 5mM MgCl2, 10mM Tris 7.5, 1mM 2-Mercaptoethanol 1X cOmplete protease inhibitor, 0.5U/ul of RNAse In Plus (Promega)). 4 mL of 29% iodoxanol cushion was prepared with a Diluent medium (250 mM Sucrose, 150mM KCl, 30mM MgCl2, 60mM Tris 8) and was loaded into an ultracentrifuge tube. 5.3 mL of sample in homogenization buffer + gradient medium was gently layered on top of the 29% iodoxanol cushion. Sample was centrifuged at 7700 xg, 4°C for 30 minutes and the supernatant was gently removed without disturbing the nuclei pellet. Nuclei were resuspended in 100 μL of Nuclei buffer (1% BSA in PBS + 1U/ul of RNAse Inhibitor).

For the preparation of nuclei for ATAC seq, we used a slightly modified protocol to extract nuclei from snap-frozen mouse cortex samples. Briefly, a ~1 cm^3^ frozen piece of mouse cortex tissue was transferred to 1 mL of ice-cold homogenisation buffer (320 mM Sucrose, 10 mM NaCl, 3mM Mg(OAc), 10mM Tris 7.5, 0.1mM EDTA, 0.1% IGEPAL-CA360, 1X cOmplete protease inhibitor and 1 mM DTT) in a Dounce homogenizer mortar and thawed for 2 minutes. The tissue was homogenised with 10 strokes of pestle A and 5 strokes of pestle B until a homogeneous nuclei suspension was achieved. The resulting homogenate was filtered through a 70 μm cell strainer (Corning). 2.65 mL of ice-cold gradient medium was added to 2.65 mL of homogenate and mixed well. 4 mL of 29% iodoxanol cushion (129.2 mM Sucrose, 77.5 mM KCl, 15.5 mM MgCl, 31 mM Tris-HCl pH 7.5, 29% Iodoxanol) was loaded into ultracentrifuge tube. 5.3 mL of sample in homogenization buffer + gradient medium was gently layered on top of the 29% iodoxanol cushion. Sample was centrifuged at 7700 xg, 4°C for 30 minutes and the supernatant was gently removed without disturbing the nuclei pellet. Nuclei were resuspended in 100 μL of Nuclei buffer (1% BSA in PBS). For the HyDrop-ATAC experiment, quadrant X was used. For the HyDrop-RNA experiment, quadrant X was used.

### HyDrop-ATAC library preparation

50 000 nuclei were resuspended in 50 μL of ATAC Reaction Mix (10% DMF, 10% Tris-HCl pH 7.4, 5 mM MgCl2, 5 ng/μL Tn5, 70 μM Pitstop, 0.1% Tween-20, 0.01% Digitonin) and incubated at 37 °C for 1 hour. 100 μL of 0.1% BSA in PBS was added and the nuclei were pelleted at 500 xg, 4 °C for 5 minutes and resuspended in 40 μL of 0.1% BSA in PBS.

Tagmented nuclei were added to 100 μL of PCR mix (1.3X Phusion HF Buffer, 15% Optiprep, 1.3 mM dNTPs, 39 mM DTT, 0.065 U/μL Phusion HF Polymerase, 0.065 U/μL Deep Vent Polymerase, 0.013 U/μL ET SSB). PCR mix was co-encapsulated with 35 μL of freshly thawed HyDrop-ATAC beads in HFE-7500 Novac oil with EA-008 surfactant (RAN Biotech) on the Onyx microfluidics platform (Droplet Genomics). The resulting emulsion was collected in aliquots of 50 μL total volume and thermocycled according to the PCR1 program (72 °C 15 min., 98 °C 3 min., 13 PCR cycles of [98 °C 10 s, 63 °C 30 s, 72 °C 1 min.], followed by a final hold on 4 °C). 125 μL of recovery agent (20% PFO in HFE), 55 μL of GITC Buffer (5 M GITC, 25 mM EDTA, 50 mM Tris-HCl pH 7.4) and 5 μL of 1 M DTT was added to each separate aliquot of 50 μL thermocycled emulsion and incubated on ice for 5 minutes. 5 μL of Dynabeads was added to the aqueous phase and incubated for 10 minutes. Dynabeads were pelleted on a Nd magnet and washed twice with 80% EtOH. Elution was performed in 50 μL of EB-DTT-Tween (10 mM DTT, 0.1% Tween-20 in EB (10 mM Tris-HCl, pH 8.5)). A 1x Ampure bead purification was performed according to manufacturer’s recommendations. Elution was performed in 30 μL of EB-DTT (10 mM DTT in EB). Eluted library was further amplified in 100 μL of PCR2 mix (1x KAPA HiFi, 1 μM index i7 primer, 1 μM index i5 primer). Final library was purified in a 0.4x-1.2x double-sided Ampure purification and eluted in 25 μL of EB-DTT (10 mM DTT in EB).

### HyDrop-RNA single cell library preparation

For a recovery of 2000 cells, 3795 cells were resuspended in 85 μL of RT mix (1x Maxima RT Buffer, 0.9 mM dNTPs, 25 mM DTT, 1.3 mM GTP, 15 % Optiprep, 1.3 U/μL RNAse inhibitor, 15 U/μL Maxima hRT, 12.5 μM TSO, 4.4% PEG-8000). RT mix was co-encapsulated with 35 μL of freshly thawed HyDrop-RNA beads in RAN oil on the Onyx microfluidics platform. The resulting emulsion was collected in aliquots of 50 μL total volume and thermocycled according to the RT program (42 °C for 90 min., 11 cycles of [50 °C for 2 min., 42 °C for 2 min.], 85 °C for 5 min., followed by a final hold on 4 °C). 125 μL of recovery agent (20% PFO in HFE), 55 μL of GITC Buffer (5 M GITC, 25 mM EDTA, 50 mM Tris-HCl pH 7.4) and 5 μL of 1 M DTT was added to each separate aliquot of 50 μL thermocycled emulsion and incubated on ice for 5 minutes. 99 μL of Ampure XP beads was added to the aqueous phase and incubated for 10 minutes. Ampure beads were pelleted on a Nd magnet and washed twice with 80% EtOH. Elution was performed in 30 μL of EB-DTT-Tween (10 mM DTT, 0.1% Tween-20 in EB). Exonuclease treatment was performed by adding 4 μL of 10x NEBuffer 3.1, 4 μL of Exo I, and 2 μL of dH2O to 30 μL of eluted library. The Exo I reaction mix was incubated at 37 °C for 5 min., 80 °C for 1 min. for heat inactivation. followed by a final hold at 4 °C. 2 μL of 1 M DTT was added and a 0.8x Ampure XP purification was performed according to manufacturer’s recommendations. cDNA was eluted in 40.5 μL of EB-DTT (10 mM DTT in EB) and added to ISPCR mix (40 μL library, 50 μL 2x KAPA HiFi, 10 μL 10 μM TSO-P primer). PCR cycling was performed according to the ISPCR program (95 °C for 3 min., 13 cycles of [98 °C for 20s, 63 °C for 20s, 72 °C for 3 min.], 72 °C for 5 min. followed by a final hold at 4 °C. 2 μL of 1M DTT was added and a 0.6x Ampure XP purification was performed according to manufacturer’s recommendations. cDNA was eluted in 28.5 μL of EB-DTT. Final sequencing library was prepared according to the following customised NEB Ultra II FS protocol (NEB E7805S). 80 ng of amplified cDNA was fragmented in Ultra II fragmentation mix (26 μL of amplified cDNA, 7 μL of NEBNext Ultra II FS Reaction Buffer, 2 μL of NEBNext Ultra II FS Enzyme Mix) on the following thermocycling program: 37 °C for 10 min., 65 °C for 30 min. and a final hold at 4 °C. 15 μL of EB was added and a 0.8x Ampure purification was performed according to manufacturer’s recommendation and eluted in 35 μL. Fragmented library was adapter-ligated in NEBNext Ultra II adapter ligation mix (35 μL of fragmented library, 30 μL of NEBNext Ultra II Ligation Master Mix, 1 μL of NEBNext Ligation Enhancer, 2.5 μL of NEBNext Adapter for Illumina) at 20 °C for 15 min., with 4 °C final hold. 28.5 μL of EB was added and a 0.8X Ampure purification was performed according to manufacturer’s recommendation and eluted in 30 μL. Eluted library was amplified in PCR master mix (50 μL 2x KAPA HiFi, 10 μL 10 μM Hy-i7 primer, 10 μL 10 μM Hy-i5 primer, 30 μL eluted library) in the following thermocycling program: 95 °C for 3 min., 13 cycles of [98 °C for 20 s, 64 °C for 30 s, 72 °C for 30 s], 72 °C for 5 min. and a final hold at 4 °C. Sequencing-ready library was purified using a 0.8x Ampure purification and eluted in 30 μL of EB.

### HyDrop-RNA optimisation trials

We performed 6 trials on a 50:50 mixture of human melanoma (MM087) and mouse melanoma (MMel). Trials were performed as described in the general HyDrop-RNA protocol, but with the following changes. All trials, except for the GTP/PEG trial, were performed using the following RT reaction mix (1.6x Maxima h-RT buffer, 1.6 mM dNTPs, 47 mM DTT, 15% Optiprep, 1.6 U/μL RNAse Inhibitor, 15.7 U/μL Maxima hRT, 12.5 μM TSO). For the Exo-condition, the Exonuclease I treatment was skipped. For all other conditions the Exonuclease I treatment was performed as described above. For the TSO-LNA trial, a locked nucleic acid TSO was used instead of the regular TSO. For the GTP/PEG trial, all steps were performed as described in the main protocol.

For the Klenow fragment second strand synthesis trial, the purified first strand product was treated with 1 μL of E. coli RNase H (NEB M0297S). The mixture was incubated at 37 °C for 30 minutes after which the enzyme was inactivated using 10 mM EDTA. The single stranded product was purified using 1.2x Ampure XP bead purification (BD sciences) and eluted in 25 μL of EB buffer. dN-SMRT primer was added to the single strand product to a final concentration of 2.5 μM and the mixture was denatured by incubation at 95 °C for 5 minutes. The sample was then allowed to cool to room temperature and incorporated in the Klenow enzyme mix (1x Maxima h-RT buffer, 1mM dNTP, 1U/μL of Klenow Exo-; NEB M0212L) was added to the single strand library. The Klenow enzyme mix was incubated at 37 °C for 60 min. The second strand reaction was stopped by heating the product at 85 °C for 5 min. The sample was purified using 1X Ampure XP and eluted in 40 μL of EB buffer. The purified second strand product was amplified with ISPCR primers as described above.

For the BST 2.0 polymerase second strand synthesis trial, the purified first strand product was treated with 1 μL of E. coli RNase H (NEB M0297S). The mixture was incubated at 37 °C for 30 minutes after which the enzyme was inactivated using 10 mM EDTA. The single stranded product was purified using 1.2X Ampure XP bead purification (BD sciences) and eluted in 25 μL of EB buffer. dN-SMRT primer was added to the single strand product to a final concentration of 2.5 μM and the mixture was denatured by incubation at 95 °C for 5 minutes. The sample was then allowed to cool to room temperature and incorporated in the Bst 2.0 enzyme mix (1X Isothermal amplification buffer, 1mM dNTP, 1U/μL of Bst 2.0 DNA polymerase; NEB M0537L) was added to the denatured library and the mixture was incubated at 55 °C for 10 mins and 60 °C for 45 minutes. The second strand reaction was stopped by heating the product at 85 °C for 5 minutes. The sample was purified using 1X Ampure XP and eluted in 40 μL of EB buffer. The purified second strand product was amplified with ISPCR primers as described above.

### Sequencing

HyDrop-ATAC libraries were sequenced on Illumina NextSeq500 or NextSeq2000 systems using 50 cycles for read 1 (ATAC paired-end mate 1), 52 cycles for index 1 (barcode), 10 cycles for index 2 (sample index) and 50 cycles for read 2 (ATAC paired-end mate 2).

HyDrop-RNA libraries were sequenced on Illumina NextSeq2000 systems using 50 cycles for read 1 (3’ cDNA), 10 cycles for index 1 (sample index, custom i7 read primer), 10 cycles for index 2 (sample index) and 58 cycles for read 2 (barcode + UMI, custom read 2 primer).

### HyDrop-ATAC data processing

Barcode reads were trimmed to exclude the intersub-barcode PCR adapters using a mawk script. Next, the VSN scATAC-seq pre-processing pipeline^22^ was used to map the reads to the reference genome and generate a fragments file for downstream analysis. Here, barcode reads were compared to a whitelist (of 884736 valid barcodes), and corrected, allowing for a maximum 1 bp mismatch. Uncorrected and corrected barcodes were appended to the fastq sequence identifier of the paired-end ATAC-seq reads. Reads were mapped to the reference genome using bwa-mem with default settings, and the barcode information was added as tags to each read in the bam file. Duplicate-marking was performed using samtools markdup. In the final step of the pipeline, fragments files were generated using Sinto (https://github.com/timoast/sinto). For mixed-species data, cells were filtered for a minimum of 1000 unique fragments and a minimum TSS enrichment of 7. For mouse cortex data, higher level analysis such as clustering and differential accessibility were performed using cisTopic^25^. In brief, cells were filtered for a minimum of 1000 unique fragments and a minimum TSS enrichment of 5. Fragments overlapping mouse candidate cis-regulatory regions^38^ were counted, and the resulting matrix was filtered for potential cell doublets using a Scrublet^24^ threshold of 0.35. Cells were Leiden-clustered based on the cell-topic probability matrix generated by an initial cisTopic LDA incorporating 51 topics, at a resolution of 0.9 with 10 neighbours. A consensus peak set was generated from per-cluster peaks and used to recount fragments. Cells were filtered using the same filtering parameters and a new model with 50 topics was trained. Cells were again Leiden-clustered based on the cell-topic probability matrix generated by the second LDA, at a resolution of 0.9 with 10 neighbours. Region accessibility was imputed based on binarised topic-region and cell-topic distributions. Gene activity was imputed based on Gini index-weighted imputed accessibility in a 10 kb up/downstream decaying window around each gene including promoters. Leiden clusters were annotated based on imputed gene accessibility around marker genes^27,28^. Differentially accessible regions were called using one-versus-all Wilcoxon rank-sum tests for each cell type, with an adjusted p-value of 0.05 and log2FC of 1.5. RPGC-normalized aggregate genome coverage bigwigs were generated from BAM files using DeepTools^39^. Per-cluster genome coverage tracks were generated using pyBigWig.

### HyDrop-RNA data processing

Barcode reads were trimmed to exclude the intersub-barcode PCR adapters using a mawk script. Reads were then mapped and cell-demultiplexed using STARsolo^40^ in CB_UMI_Complex mode. The resulting STARsolo-filtered count matrices were further analysed using Scanpy^41^. In short, cells were filtered on expression of a maximum of 4000 genes, and a maximum of 1% UMIs from mitochondrial genes. Genes were filtered on expression in a minimum of 3 cells. Potential cell doublets were filtered out using a Scrublet^24^ threshold of 0.25. The filtered expression matrix was scaled to total counts and log-normalized. Total counts and mitochondrial reads were regressed out and UMAP embedding was performed after PCA. Cells were annotated and fine tuned based on differential gene expression of marker genes sourced from either the Davie et al. Drosophila brain atlas^1^ or the Allen Brain RNA-seq Database^27^.

Public inDrop and Drop-seq data^5,6^ were downloaded from their respective GEO repositories. Both Drop-seq and inDrop cells were filtered for a minimum of 500 genes per cell as described in their respective papers. Public reference 10x single-cell ATAC-seq data was sourced from https://support.10xgenomics.com/single-cell-atac/datasets (“<u>Flash frozen cortex, hippocampus, and ventricular zone from embryonic mouse brain (E18)</u>”, “<u>Fresh cortex from adult mouse brain (P50)</u>”). Public reference 10x single-cell gene expression data was sourced from https://support.10xgenomics.com/single-cell-gene-expression/datasets (“<u>1k Brain Nuclei from an E18 Mouse</u>”, “<u>2k Brain Nuclei from an Adult Mouse (>8 weeks)</u>”). Public PC-3 and MCF-7 ATAC-seq data was sourced from ENCODE (ENCFF772EFK, ENCFF024FNF).

Data was visualised using a combination of Seaborn^42^ and Matplotlib^43^. A vector image representing mouse head and cortex was sourced from SciDraw^44^.

## Acknowledgements

We thank Andrew Adey for his great advice on Tn5 and scATAC-seq. We also thank Sebastián Najle, Céline Vallot, and Arnau Sebé-Pedrós for many discussions on droplet microfluidics, Frederik Ceyssens and the KU Leuven Nanocenter for their support with microfabrication, Ghanem Ghanem for his kind donation of the MM087 melanoma lines, Jean-Christophe Marine for his kind donation of the mouse melanoma lines, and Koen De Wispelaere for his assistance during his master internship.

## Ethical approval

All animal experiments were conducted according to the KU Leuven ethical guidelines and approved by the KU Leuven Ethical Committee for Animal Experimentation (approved protocol numbers ECD P037/2016, P014/2017, and P062/2017). All use of cell lines was approved by the KU Leuven Ethical Committee for Research under project number S63316.

## Author contributions

FDR, SP and SA wrote the manuscript. SP designed and fabricated the chips used for HyDrop bead production and single cell encapsulation. SP, FDR, SA conceived the HyDrop protocols and designed the experiments. SP conceived and developed the HyDrop bead barcoding strategy and optimized the protocols with FDR, KT. FDR, SP, JI performed the HyDrop experiments. GM performed mouse cortex microdissections. CF, GH, FDR developed the HyDrop pre-processing and mapping pipeline. FDR, CBG, JJ performed the downstream data analysis.

## Data availability statement

The datasets generated during and/or analysed during the current study are available on the Gene Expression Omnibus (GEO) with the primary accession number GSE175684 (https://www.ncbi.nlm.nih.gov/geo/query/acc.cgi?acc=GSE175684), and on SCope (https://scope.aertslab.org/#/HyDrop/*/welcome). Step-by-step user protocols are available on Protocols.io (https://www.protocols.io/workspaces/aertslab). Data analysis tutorials for HyDrop are available on GitHub (https://github.com/aertslab/hydrop_data_analysis).

