## Supplemental information for "HyDrop: droplet-based scATAC-seq and scRNA-seq using dissolvable hydrogel beads"

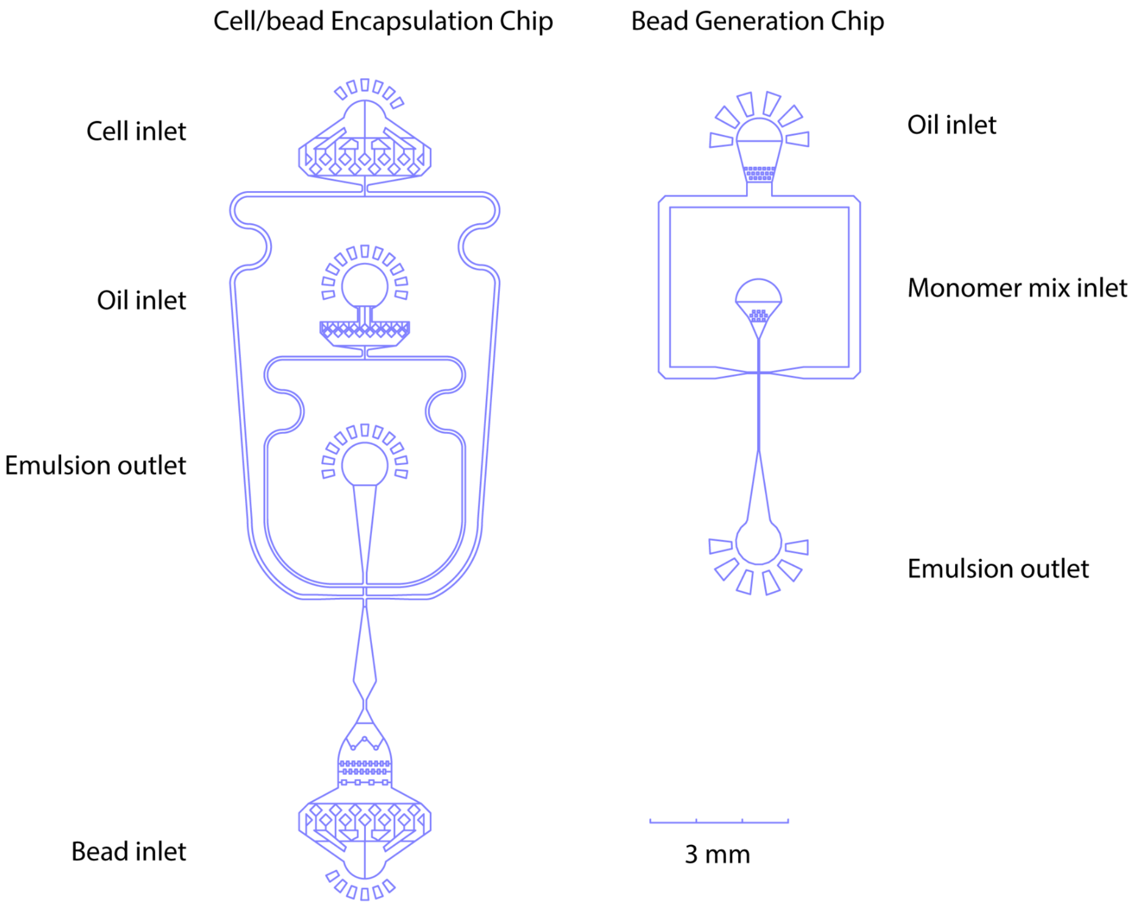


**Figure S1.** Microfluidic chip designs for cell/bead encapsulation and hydrogel bead generation. Oil, monomer mix, cell suspensions and beads can be pumped in through their respective inlets. At the flow focusing point, aequous phases (cells/beads/monomer mix) are emulsified by collision with the oil phase at the flow-focusing point. Emulsion is collected at the outlet channel.

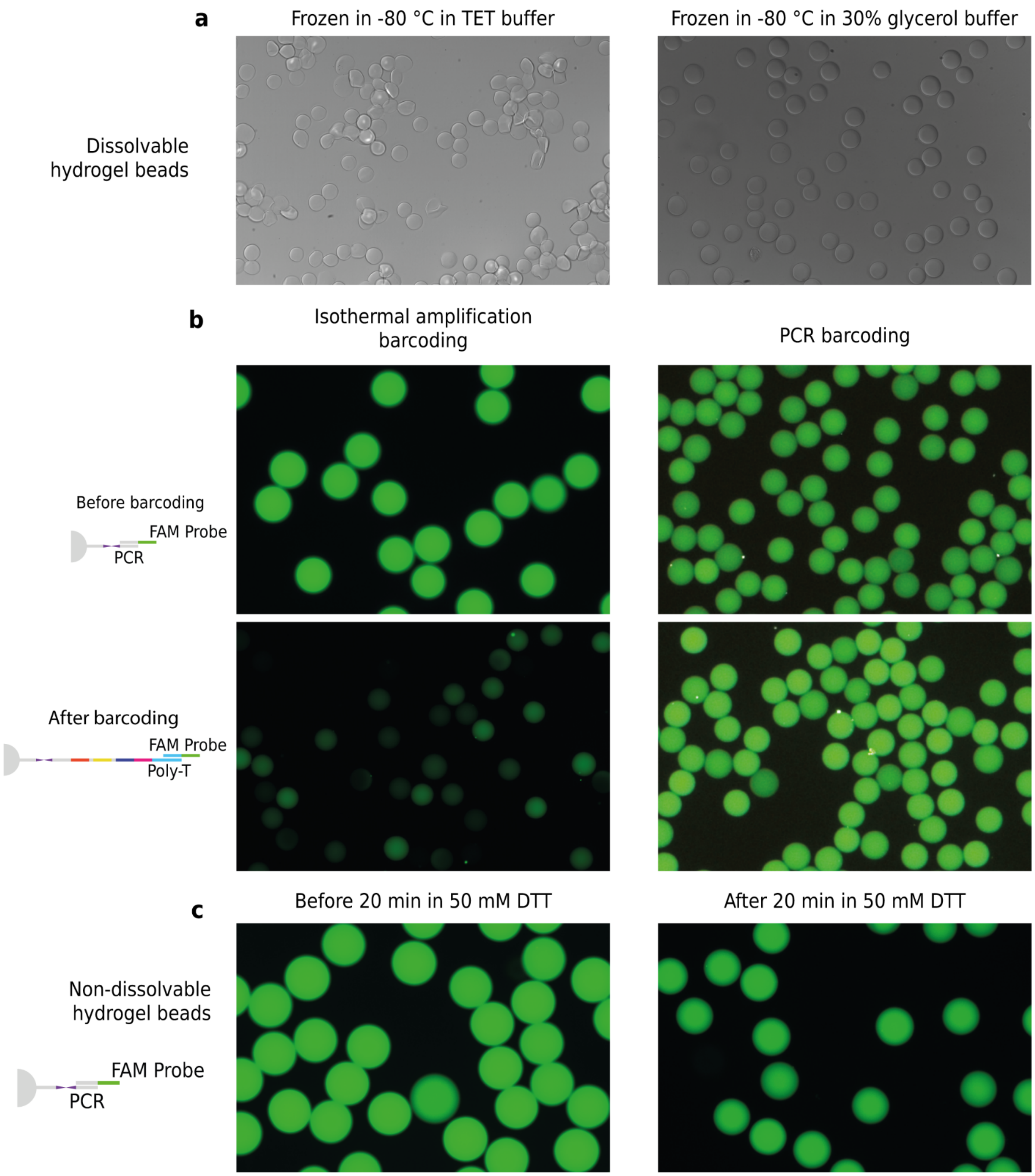


**Figure S2. a.** Hydrogel bead integrity after being frozen and thawed in TET buffer and 30% glycerol buffer. **b.** Hydrogel beads fluorescence intensity before and after barcoding for isothermal and PCR barcoding process. **c.** Non-dissolvable HyDrop-RNA beads show residual fluorescence signal even after 20 minute incubation in 50 uM of DTT.

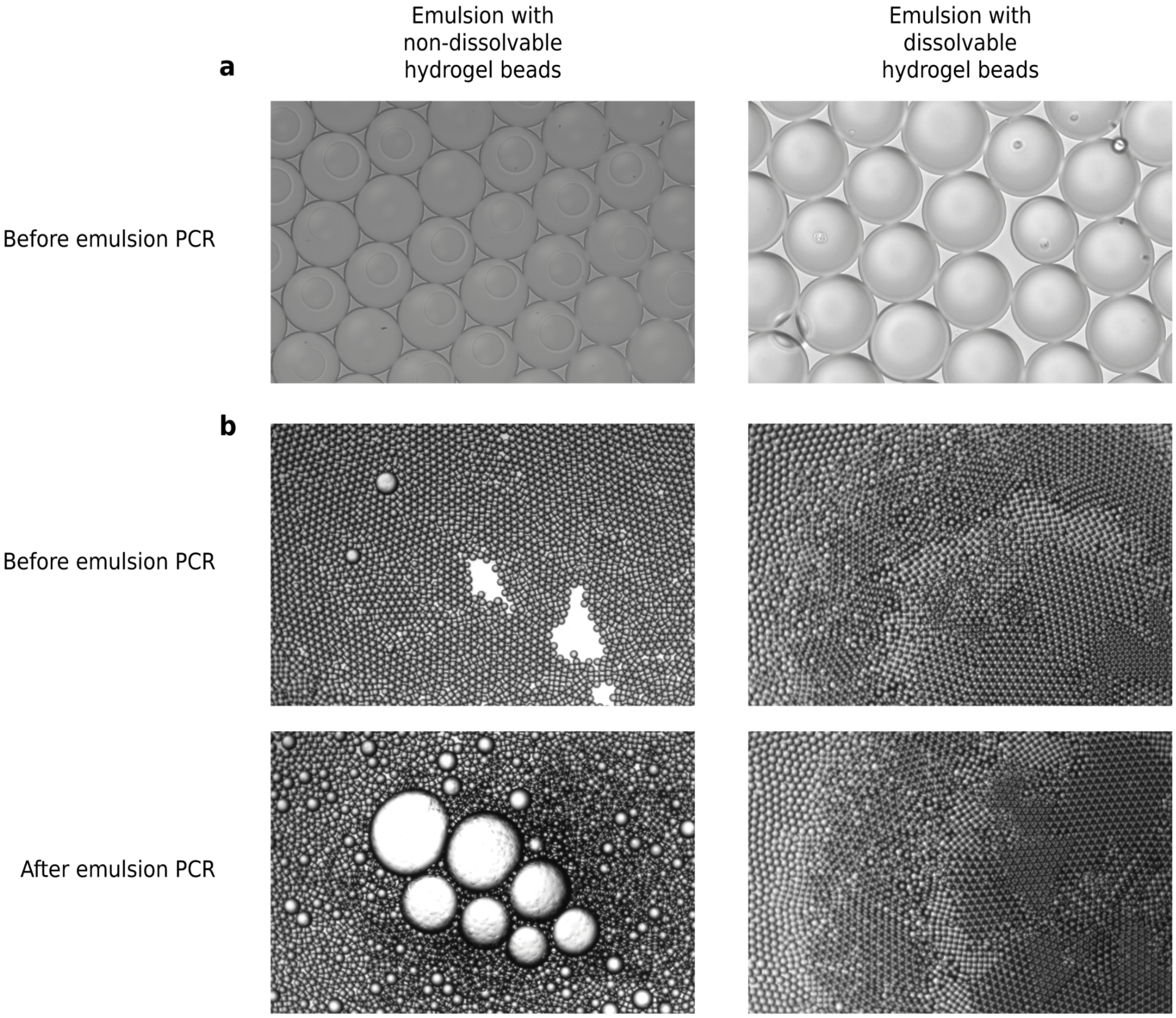


**Fig. S3**. **a.** HyDrop emulsions made with non-dissolvable and dissolvable beads, picture taken minutes after the emulsion was made showing that dissolvable beads dissolve within minutes. **b.** HyDrop-ATAC emulsions made with non-dissolvable and dissolvable beads before and after thermocycling.

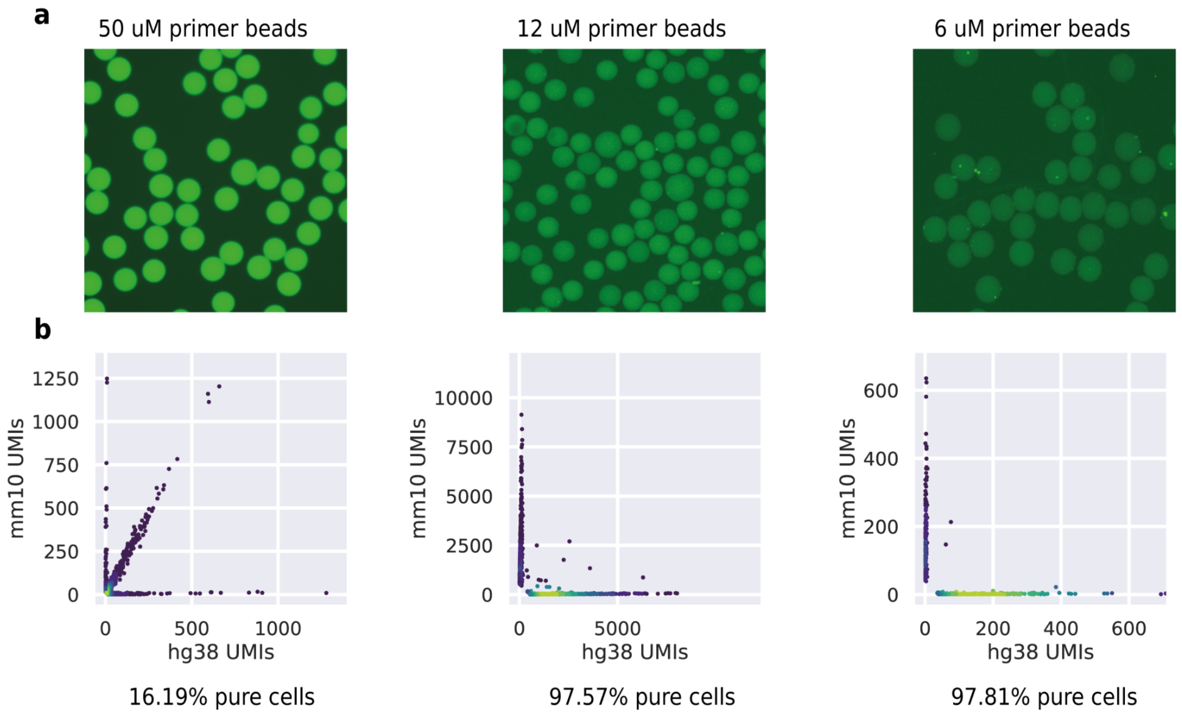


**Figure S4. a.** Fluorescence signal of 50, 12 and 6 uM primer concentration beads. **b.** HyDrop-RNA species-mixing purity plots made using 50, 12 and 6 uM beads.

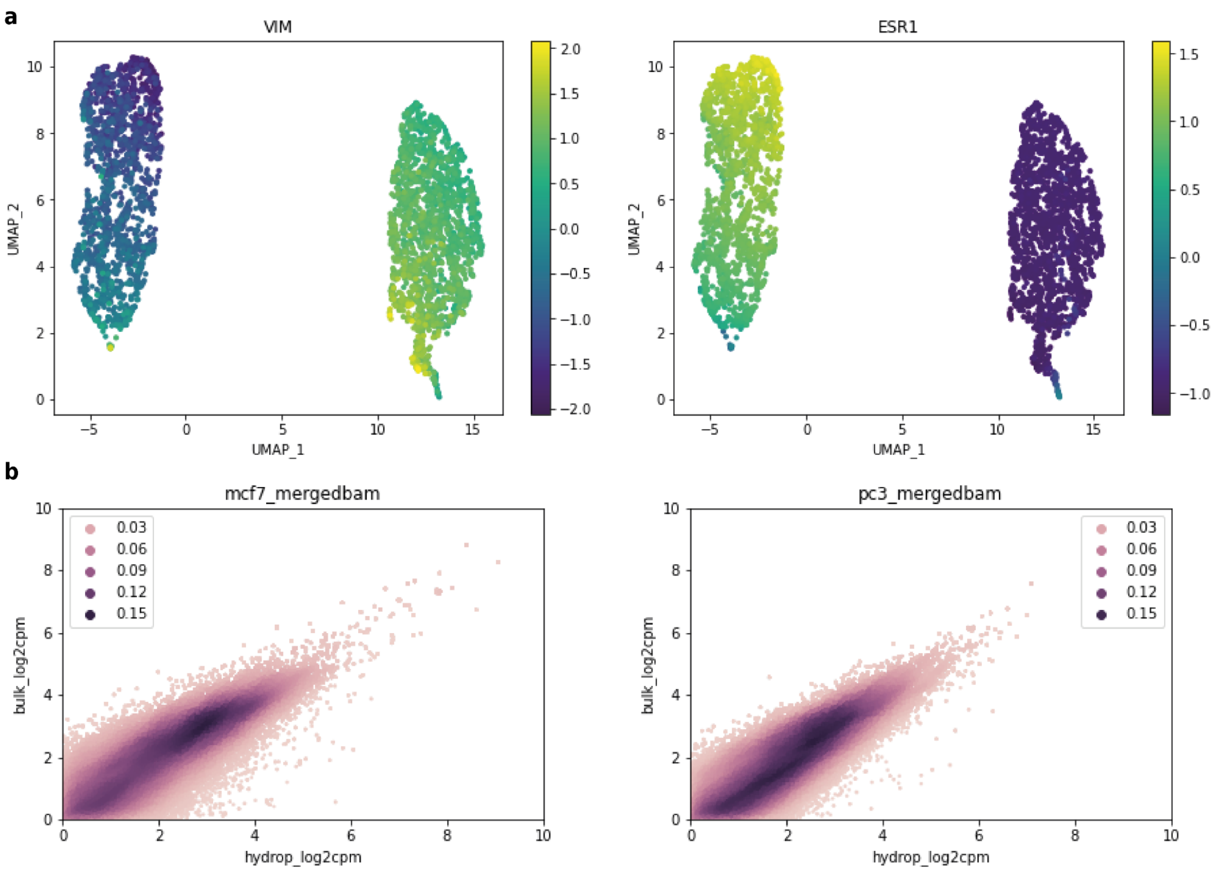


**Figure S5. a.** HyDrop-ATAC on MCF-7/PC-3 cells UMAP colored by imputed gene activity for Vimentin (VIM) and Estrogen Receptor 1 (ESR1). **b.** Correlation of HyDrop-ATAC and bulk ATAC counts in regions for MCF-7 and PC-3 cells.

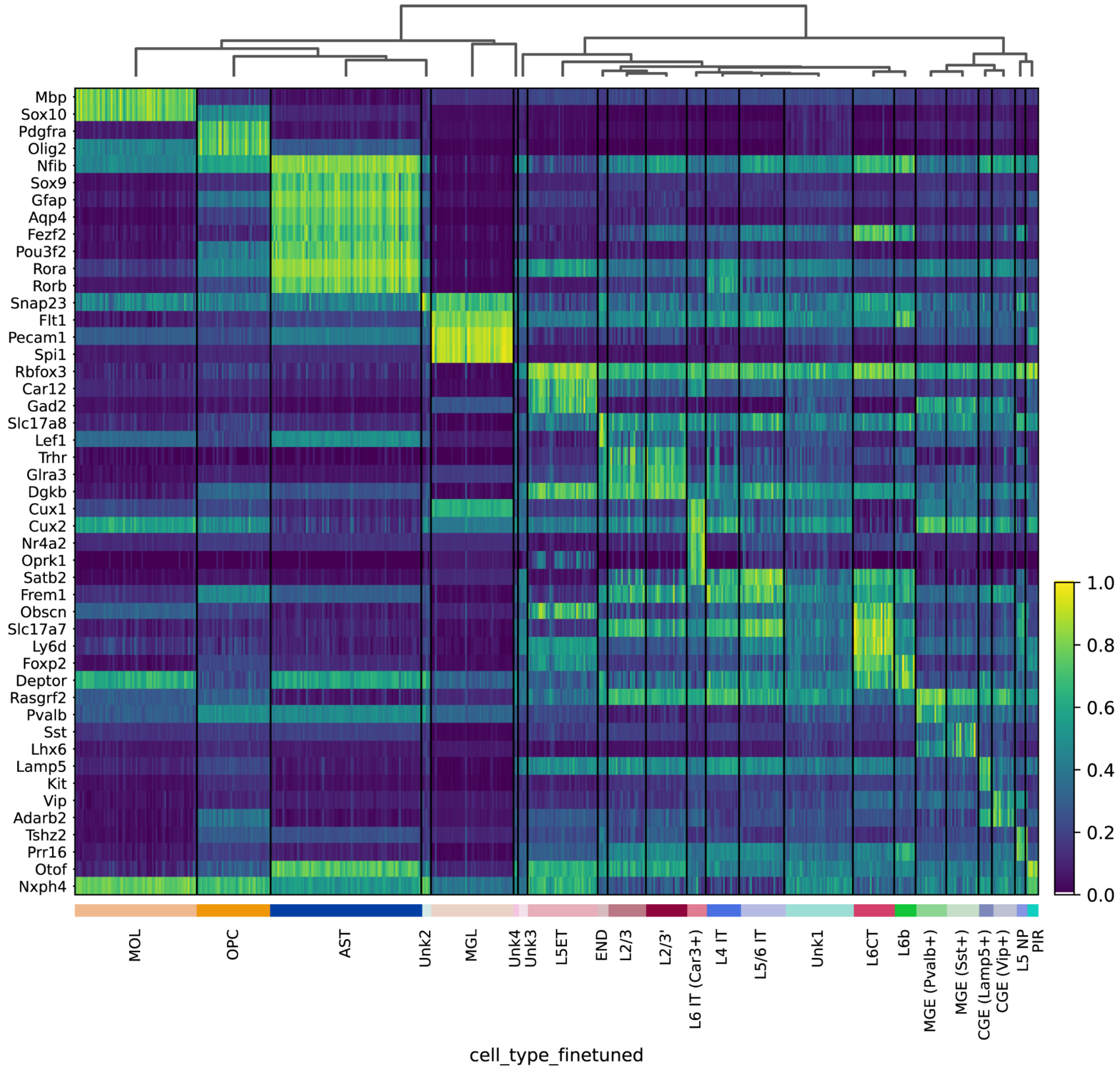


**Figure S6.** Heatmap of HyDrop-ATAC on mouse cortex gene activity imputed by accessibility within a 10 kb window around several marker genes.

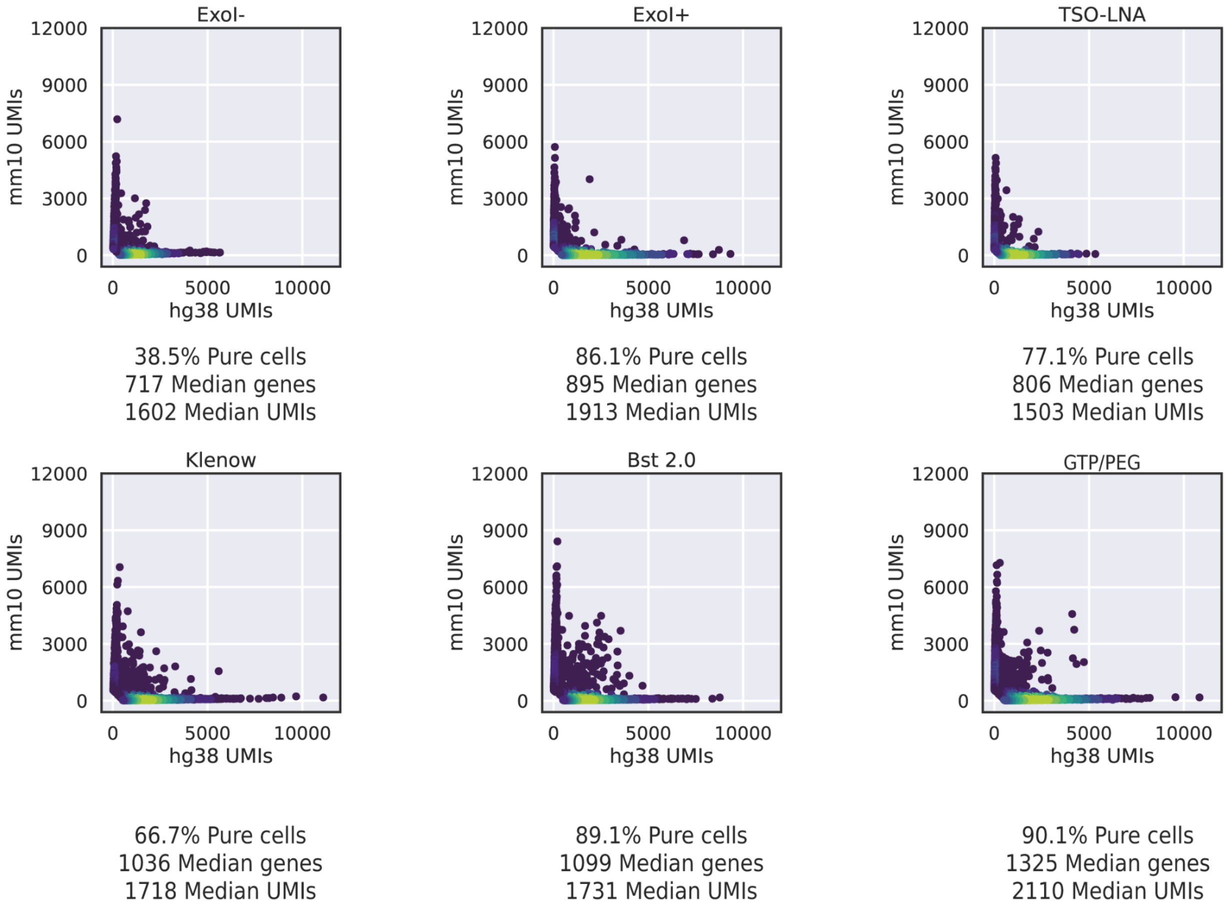


**Figure S7.** HyDrop-RNA species-mixing purity plots for Exo I treated or Exo I non-treated, TSO-LNA, GTP/PEG, Bst2.0 and Klenow fragment HyDrop-RNA libraries.

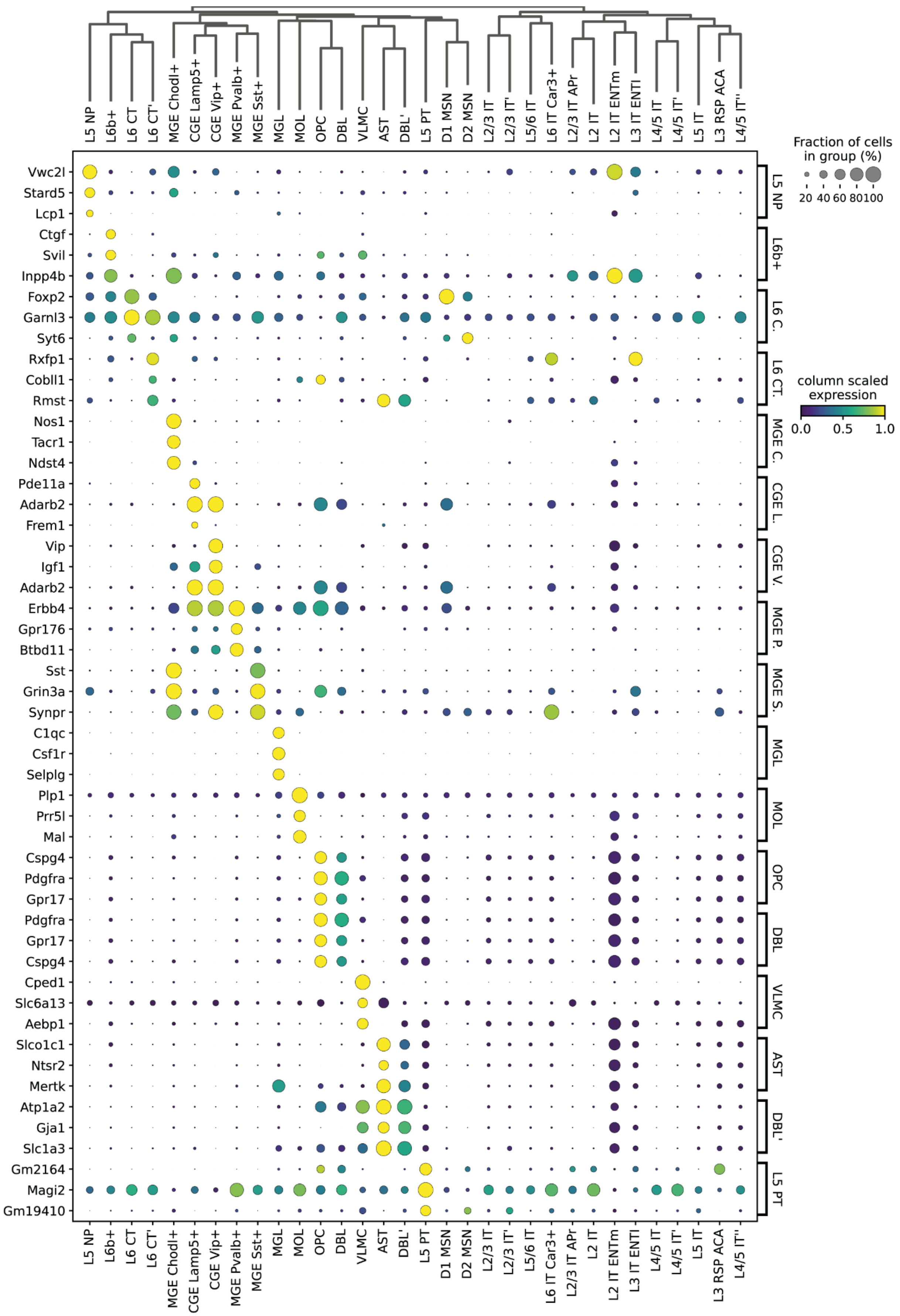

Fraction of cells in group (%)
20 40 60 80 100
column scaled expression
0.0 0.5 1.0
L5 NP
L6b+
L6 CT
L6 CT'
MGE Chodl+
CGE Lamp5+
CGE Vip+
MGE Pvalb+
MGE Sst+
MGL
MOL
OPC
DBL
VLMC
AST
DBL'
L5 PT
D1 MSN
D2 MSN
L2/3 IT
L2/3 IT'
L5/6 IT
L6 IT Car3+
L2/3 IT APr
L2 IT
L2 IT ENTm
L3 IT ENTl
L4/5 IT
L4/5 IT'
L5 IT
L3 RSP ACA
L4/5 IT''
Vwc2l
Stard5
Lcp1
Ctgf
Svil
Inpp4b
Foxp2
Garnl3
Syt6
Rxfp1
Cobll1
Rmst
Nos1
Tacr1
Ndst4
Pde11a
Adarb2
Frem1
Vip
Igf1
Adarb2
Erbb4
Gpr176
Btbd11
Sst
Grin3a
Synpr
C1qc
Csf1r
Selplg
Plp1
Prr5l
Mal
Cspg4
Pdgfra
Gpr17
Pdgfra
Gpr17
Cspg4
Cped1
Slc6a13
Aebp1
Slco1c1
Ntsr2
Mertk
Atp1a2
Gja1
Slc1a3
Gm2164
Magi2
Gm19410
L5 NP
L6b+
L6 C.
L6 CT.
MGE C.
CGE L.
CGE V.
MGE P.
MGE S.
MGL
MOL
OPC
DBL
VLMC
AST
DBL'
L5 PT

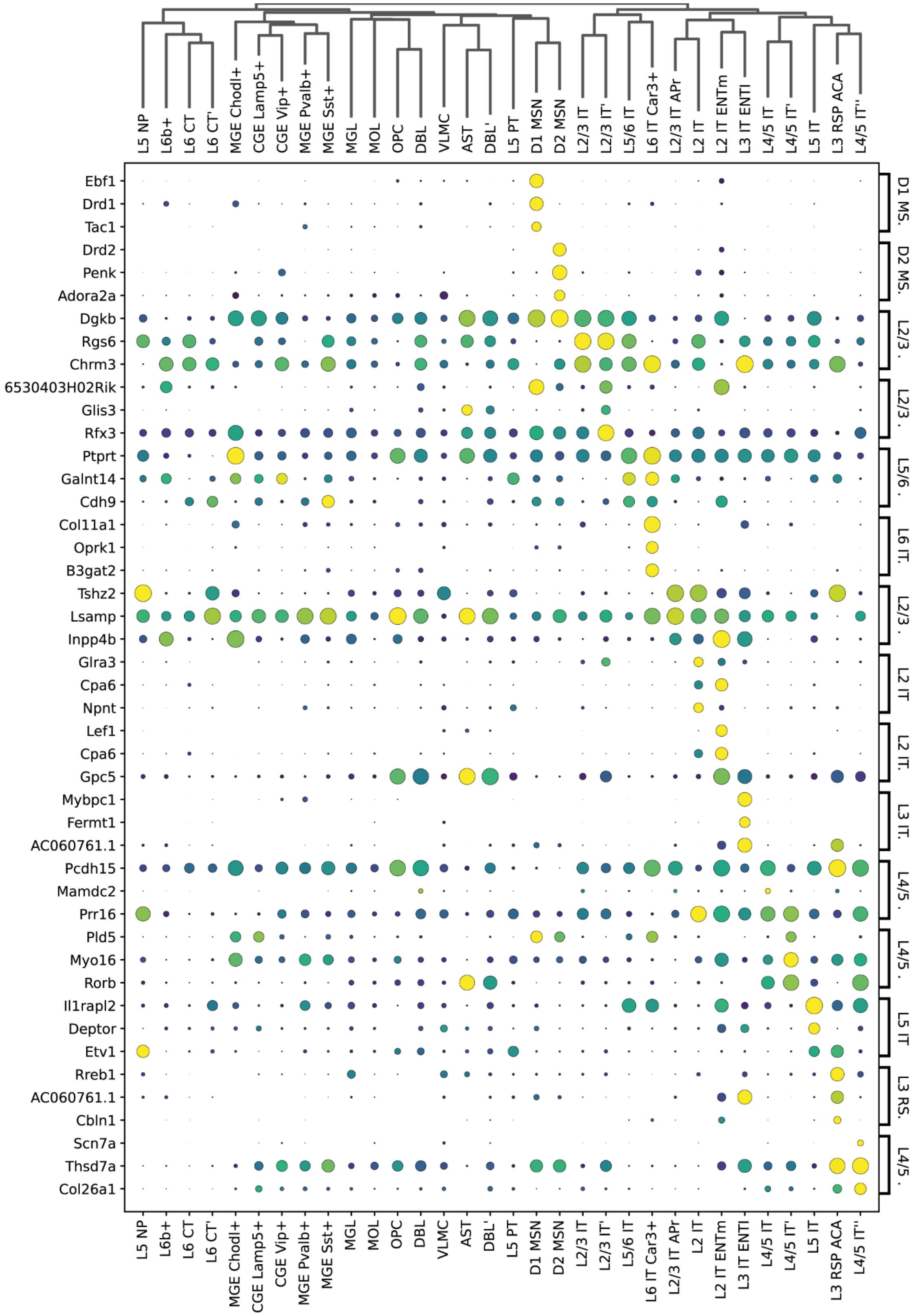


**Figure S8.** Dotplot of HyDrop-RNA mouse cortex top 3 differentially expressed genes from each cluster.

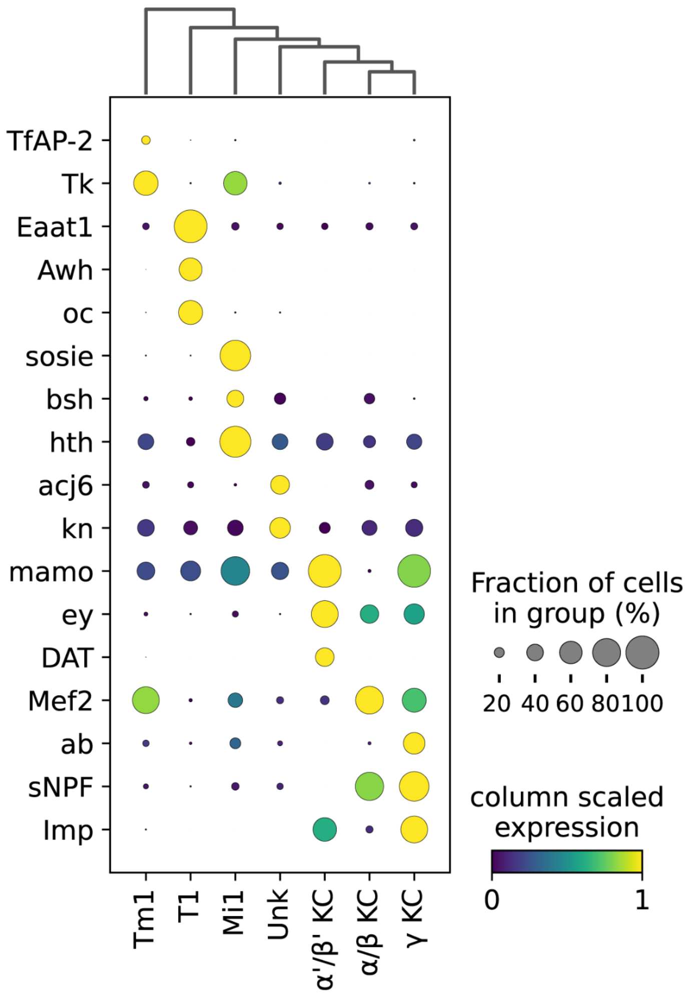


**Figure S9.** Dotplot of marker genes of HyDrop-RNA on FAC-sorted fly brain cells.
